## Supplementary Information for "FAP106 is an interaction hub required for assembly of conserved and lineage-specific microtubule inner proteins at the cilium inner junction"

### Supplemental Figures

#### Figure S1

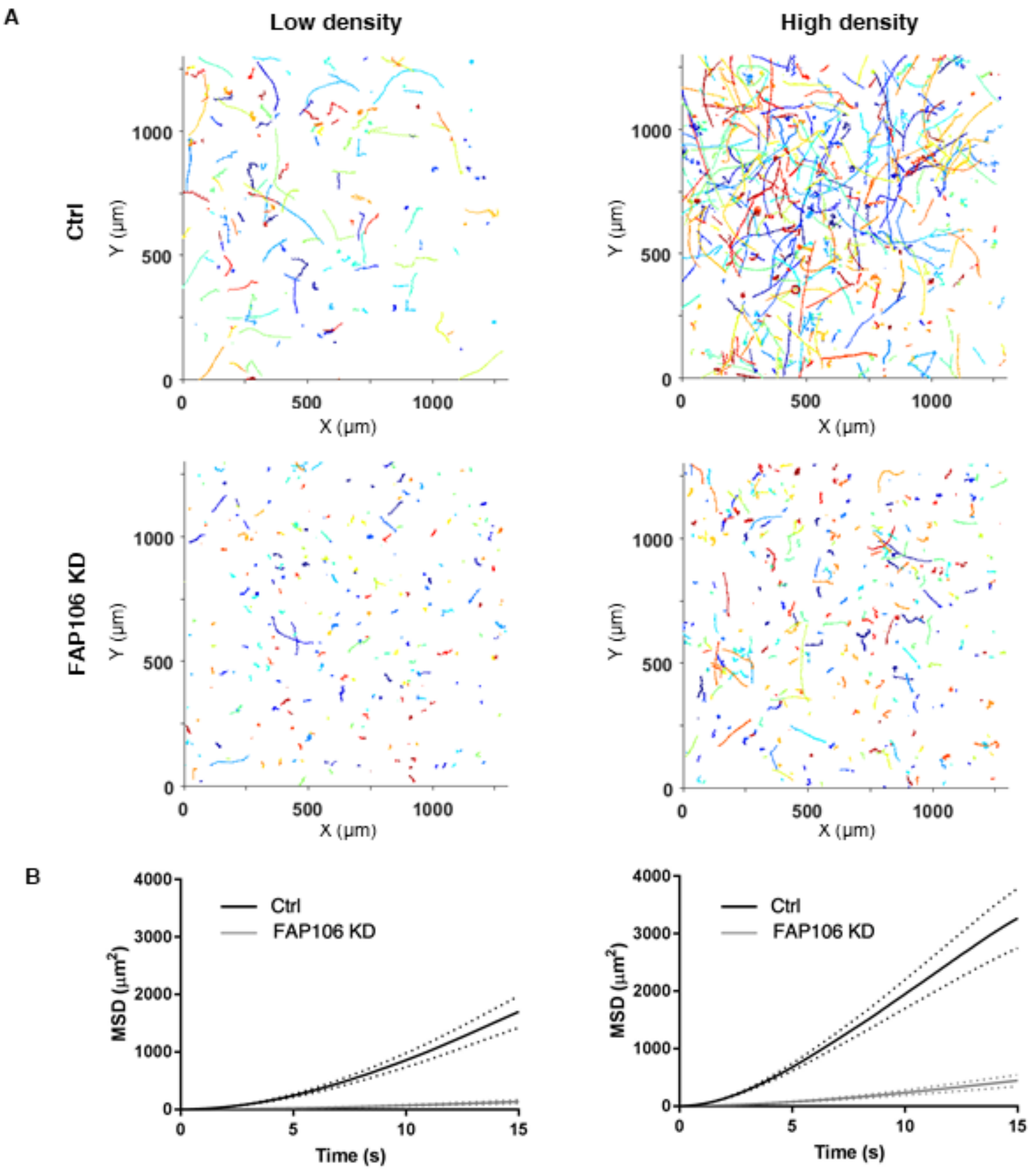


**Figure S1. Motility analysis of FAP106 KD.** (*A*) Tracks of individual 29-13 (Ctrl) or FAP106 KD parasites grown in the presence of tetracycline to induce knockdown. Parasites were grown to a density of ~1e6 cells/ml (low density) or ~1e7 cells/ml and diluted to ~1e6 cells/ml in conditioned medium just prior to imaging (high density). (*B*) Mean squared displacement of parasites tracked in *A*. Dotted lines indicate the upper and lower bounds of the standard error of the mean. Low density: Ctrl N=263, FAP106 KD N=388; High density: Ctrl N=219, FAP106 KD N=389. Data are from an independent biological experiment from Fig. 1F-G.

**Figure S2**

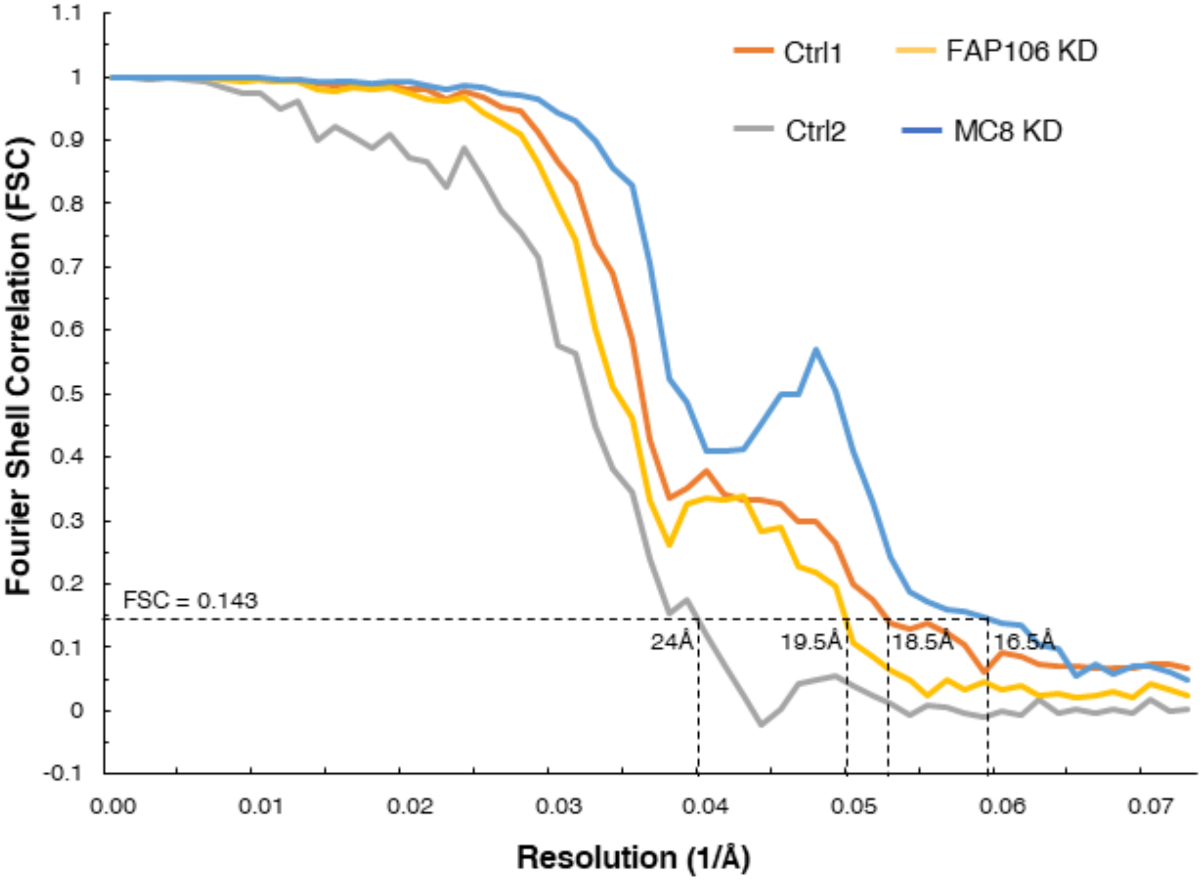


**Figure S2. Resolution evaluation of sub-tomographic averages.** FSC coefficients as a function of spatial frequency for the final sub-tomographic averages of the 48-nm repeat. Ctrl1 and Ctrl2 are from 29-13, analyzed alongside FAP106 KD (Fig. 2) and MC8 KD (Fig. 4), respectively. The effective resolution of each structure, as indicated by the dashed lines, is based on the FSC at the 0.143 coefficient cutoff.

**Figure S3**

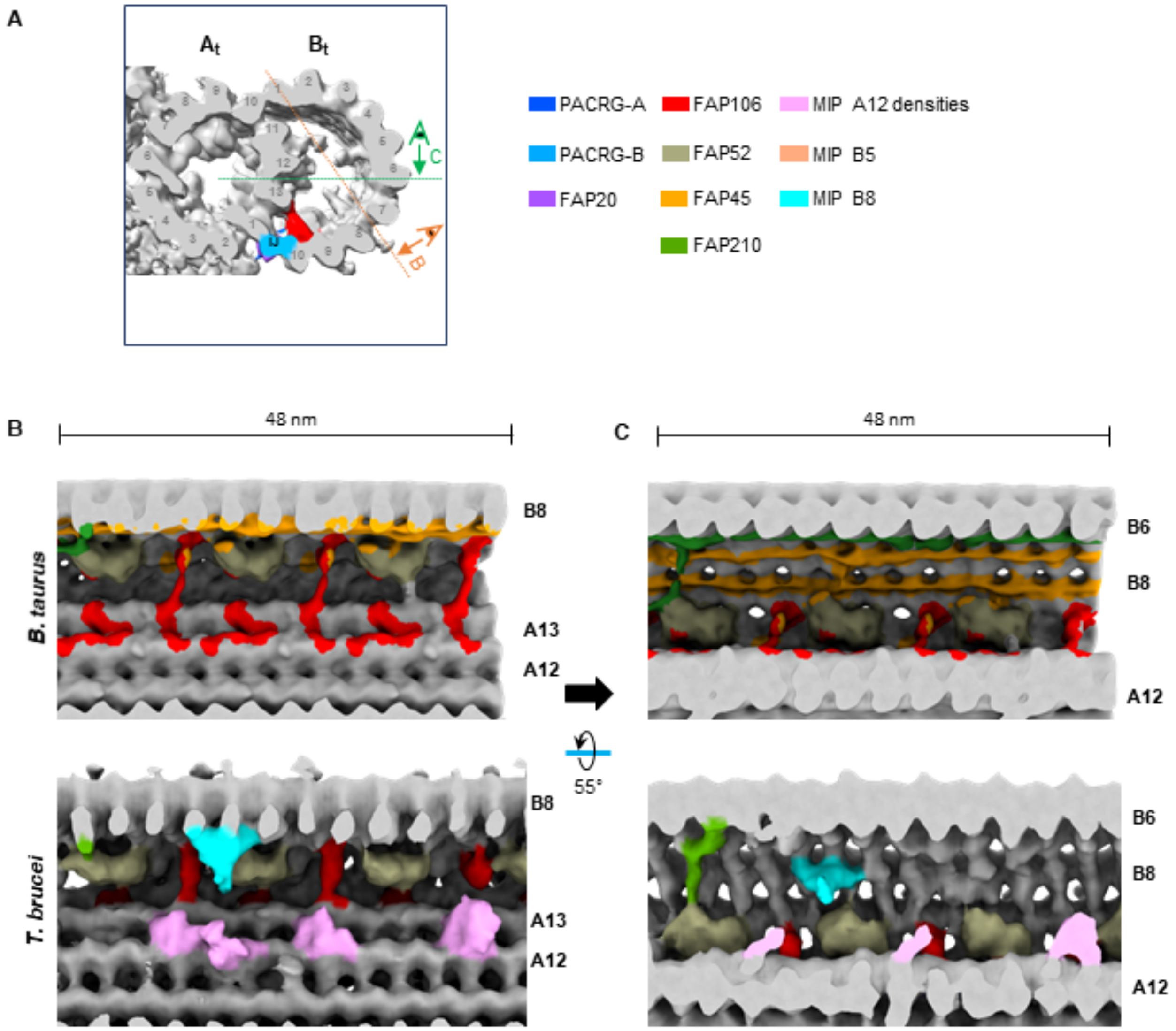


**Figure S3. Comparison of the 48-nm repeat from *T. brucei* and *B. taurus* DMTs** (*A*) Cross-sectional view of the *T. brucei* DMT indicating viewing angles shown in *B*-*C*. (*B*-*C*) Longitudinal views of the *T. brucei* DMT shown in Fig. 2 are aligned with corresponding views of the published density map from *B. taurus* (emdb-24664; (*1*)), which has been low pass filtered to a similar resolution. Conserved MIP densities are colored based on similarity to densities in *B.* taurus and other published structures (*1-4*) as indicated by the legend. Note that *T. brucei* has a structure similar to FAP210 (green) in *B. taurus* despite not having a clear FAP210 homolog (Table 1). Lineage-specific *T. brucei* densities that are not present in the *B. taurus* structure and are reduced in FAP106 KD (Fig. 2) are colored according to the legend.

**Figure S4**

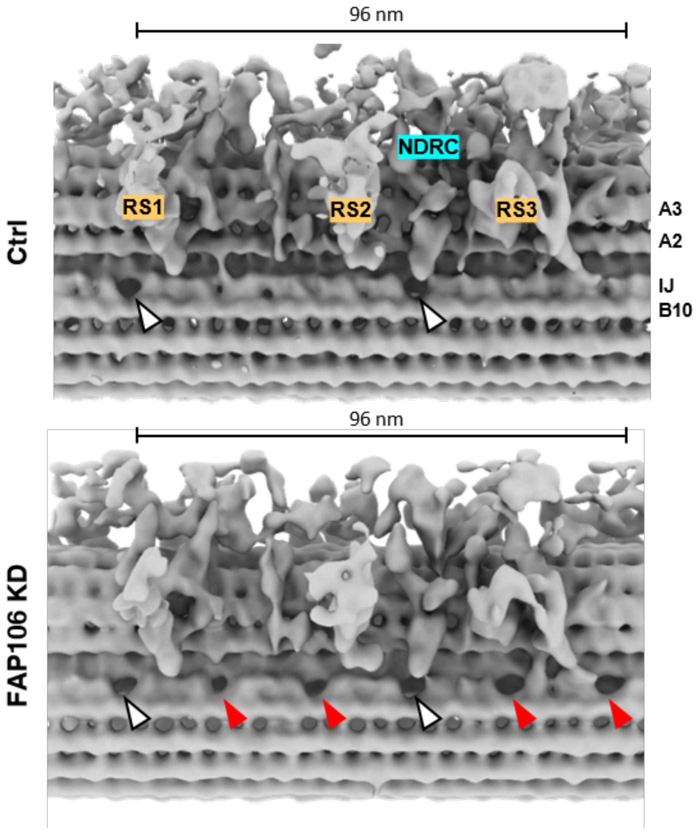


**Figure S4. Holes in the IJ filament in the 96-nm repeating unit of control and FAP106 KD DMTs.** Structure of the 96-nm repeat of the DMT was obtained by cryoET, with subtomographic averaging. Holes in the IJ filament are indicated with arrowheads. Red arrowheads indicate four additional holes in the FAP106 KD IJ filament compared to two holes in control (white arrowheads) (*5*). The nexin dynein regulatory complex (NDRC) and radial spokes (RS1-3) are labeled.

**Figure S5**

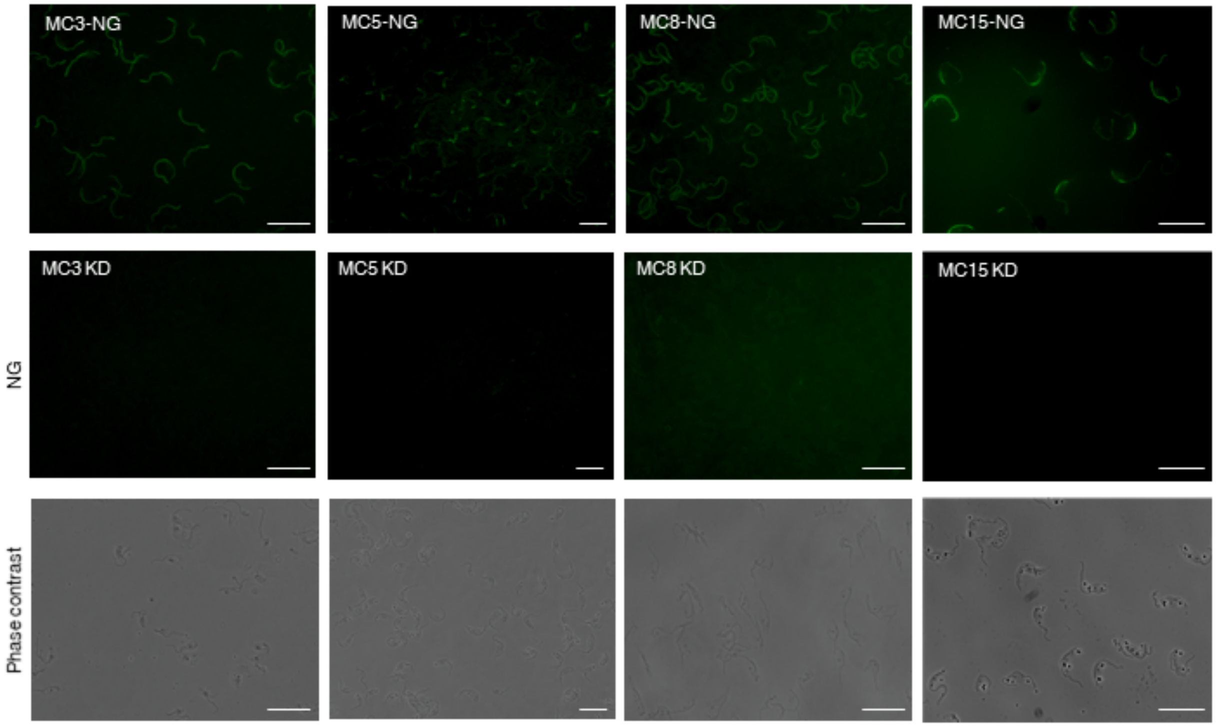


**Figure S5. Flagellar localization and knockdown of MCs.** Fluorescence microscopy of detergent-extracted cytoskeletons shows flagellar localization of mNeonGreen (NG)-tagged MCs. Knockdown (KD) constructs were introduced into the corresponding NG-tagged lines. Parasites were grown in the presence of tetracycline to induce knockdown of the indicated MCs, confirming loss of flagellar localization. Scale bar = 20 μm.

**Figure S6**

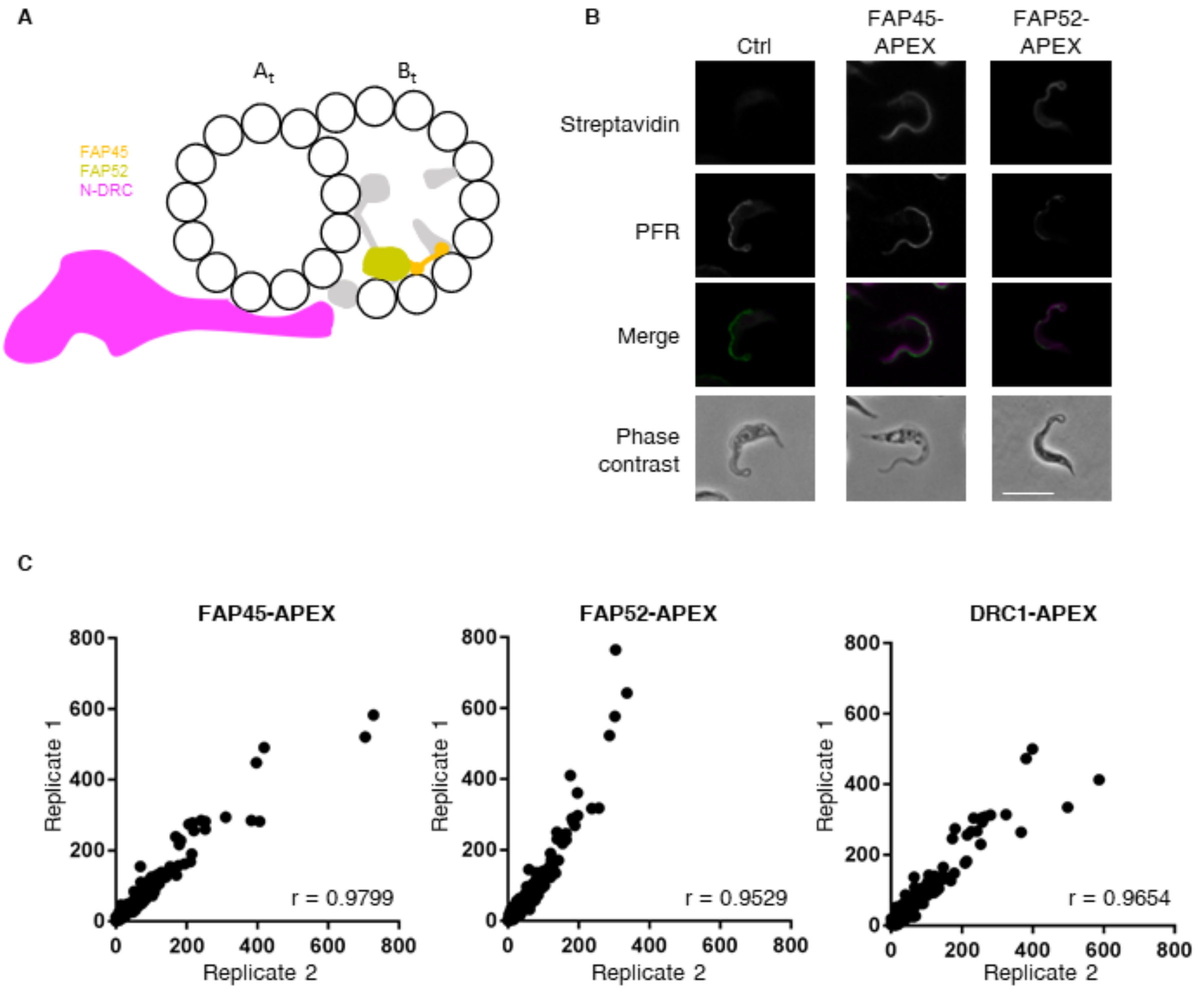


**Figure S6. APEX2-based proximity labeling of B-tubule MIPs.** (*A*) Cross-sectional view of the DMT as viewed from the flagellum tip, showing generalized view of B-tubule MIPs (grey) and indicating relative positions of three different proteins tagged with APEX2 for proximity labeling: known B-tubule MIPs FAP45 (orange) and FAP52 (tan); non-MIP control outside DMT: DRC1, part of N-DRC (pink). A- and B-tubules are labeled ($A_t$, $B_t$). (*B*) Immunofluorescence microscopy of whole cells showing APEX2-dependent biotinylation (streptavidin; magenta) in the flagellum (anti-PFR, green) of FAP45-APEX and FAP52-APEX cells. APEX2-dependent biotinylation in DRC1-APEX cells has been described previously (*6*). Scale bar = 10 um. (*C*) Plots showing correlation of protein abundance between independent biological replicates. Pearson coefficients (r) were calculated in GraphPad Prism.

**Figure S7**

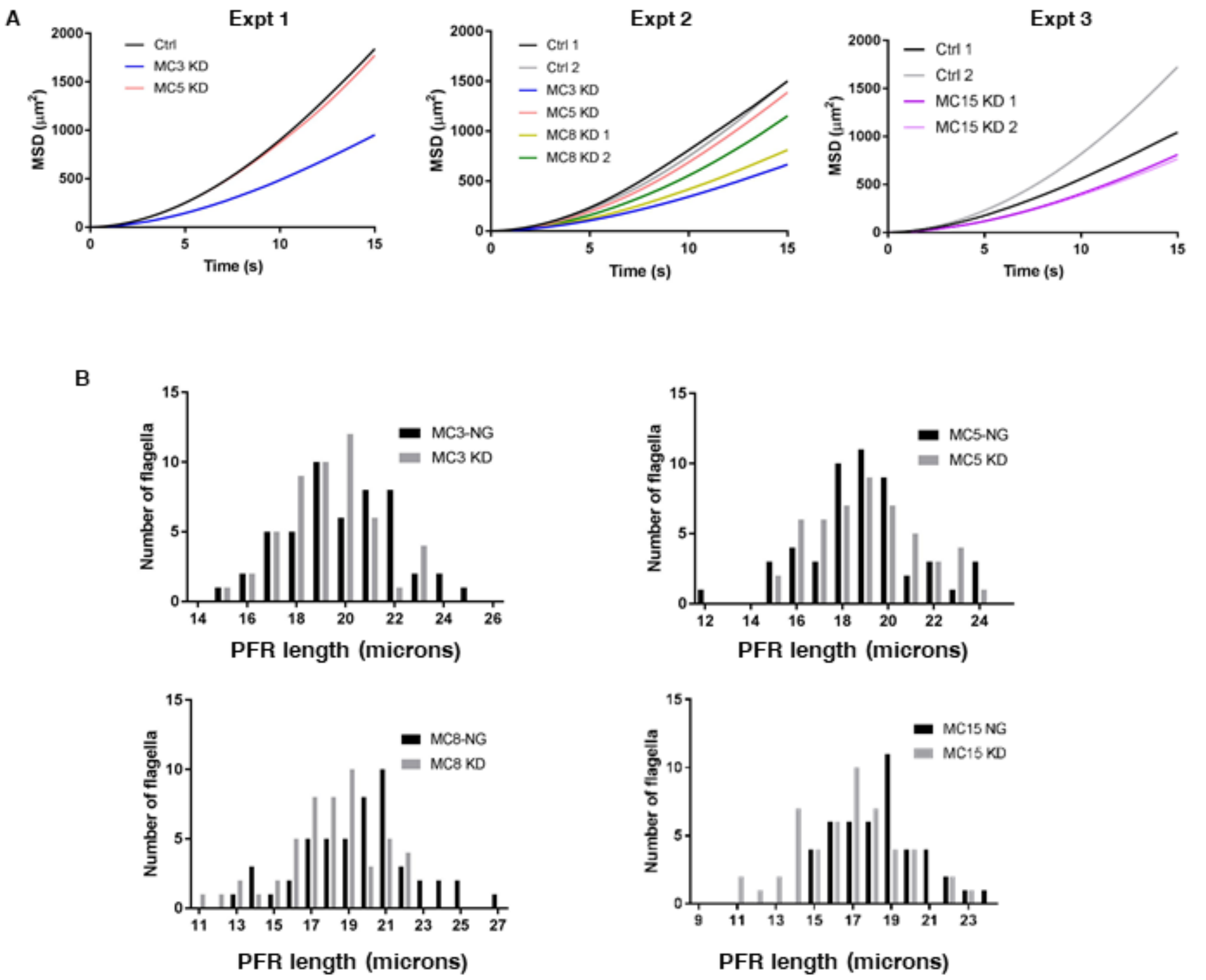


**Figure S7. Effect of MC knockdown on parasite motility and flagellum length.** Parasites were grown in the presence of tetracycline to induce knockdown of the indicated MCs. (*A*) Motility analyses showing the mean squared displacement (MSD) of parasites. Expt 1: 29-13 (Ctrl) N=361, MC3 KD N=318, MC5 KD N=415; Expt 2: 29-13 (Ctrl 1) N=309, 29-13 (Ctrl 2) N=360, MC3 KD N=309, MC5 KD N=338, MC8 KD 1 N=361, MC8 KD 2 N=348; Expt 3: 29-13 (Ctrl 1) N=342, 29-13 (Ctrl 2) N=389, MC15 KD 1 N=367, MC15 KD 2 N=384. Figure 3C shows the mean squared displacement of the three combined experiments that are shown here. (*B*) Detergent-extracted cytoskeletons were prepared from the indicated KD parasites or their respective NG-tagged parental cell lines (Ctrl). Histograms show the distribution of flagellum lengths as measured by anti-PFR immunofluorescence microscopy. Average flagellum lengths are plotted in Fig. 3D.

**Figure S8**

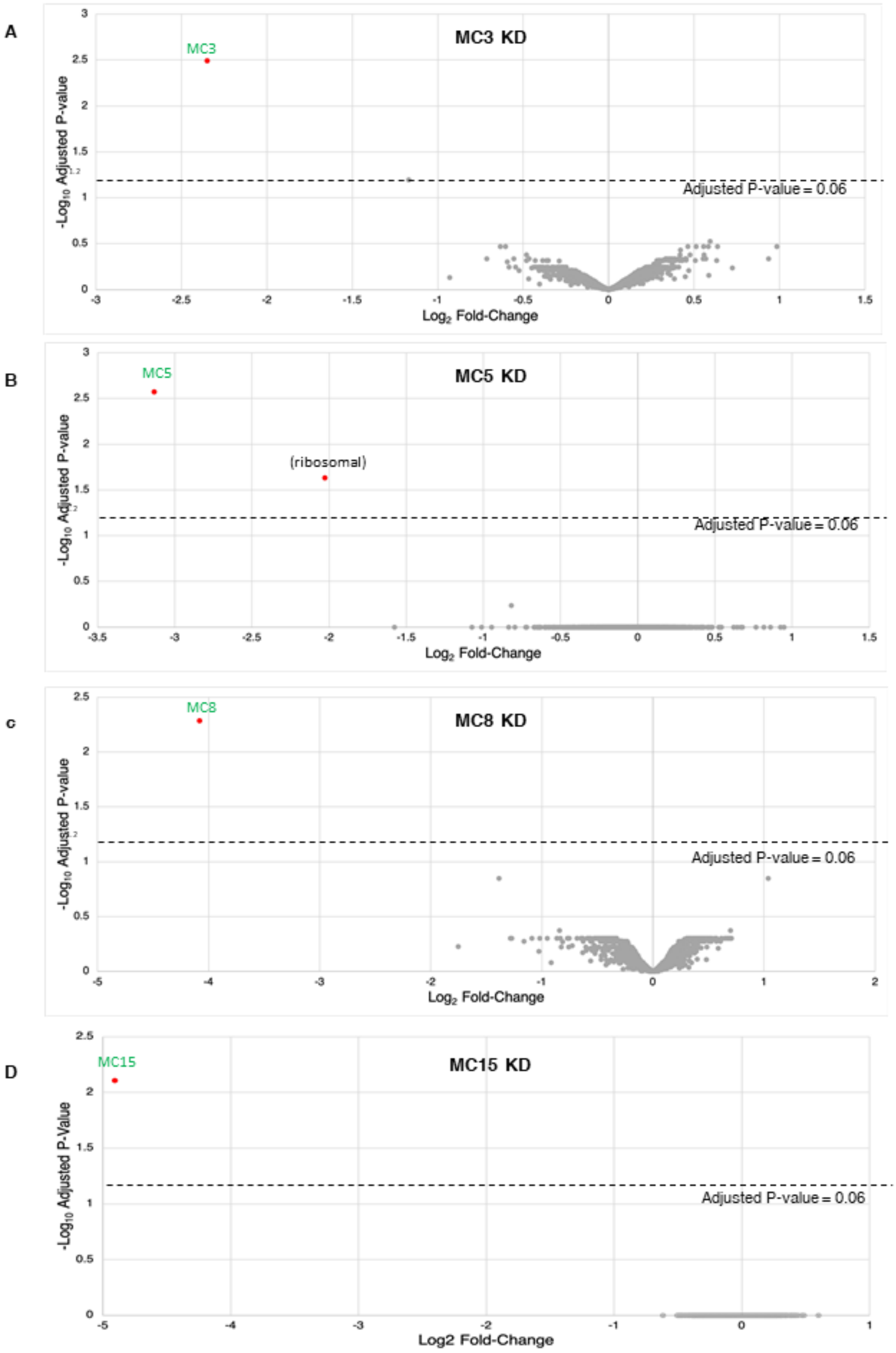
A
MC3 KD
MC3
Adjusted P-value = 0.06
-Log10 Adjusted P-value
Log2 Fold-Change
B
MC5 KD
MC5
(ribosomal)
Adjusted P-value = 0.06
-Log10 Adjusted P-value
Log2 Fold-Change
C
MC8 KD
MC8
Adjusted P-value = 0.06
-Log10 Adjusted P-value
Log2 Fold-Change
D
MC15 KD
MC15
Adjusted P-value = 0.06
-Log10 Adjusted P-Value
Log2 Fold-Change

**Figure S8. Knockdown of individual MCs affects only the targeted MC.** Parasites were grown in the presence of tetracycline to induce knockdown of the indicated MCs. Demembranated flagella were purified from the indicated KD parasites or their respective NG-tagged parental cell lines (Ctrl) and subjected to TMT quantitative proteomic analysis. Volcano plots show significance (-$Log_{10}$ Adjusted P-value) vs. relative abundance ($Log_2$ Fold-Change) for all proteins quantified by TMT proteomics. Relative abundance is calculated as MC KD/Ctrl. Results are from two independent biological samples. Proteins that met the filtering criteria (2-fold reduced, adjusted P-value $\leq$0.06; Supp. Dataset S1) are indicated by red dots. (*A*) MC3 KD. (*B*) MC5 KD. (*C*) MC8 KD. (*D*) MC15 KD.

**Figure S9**

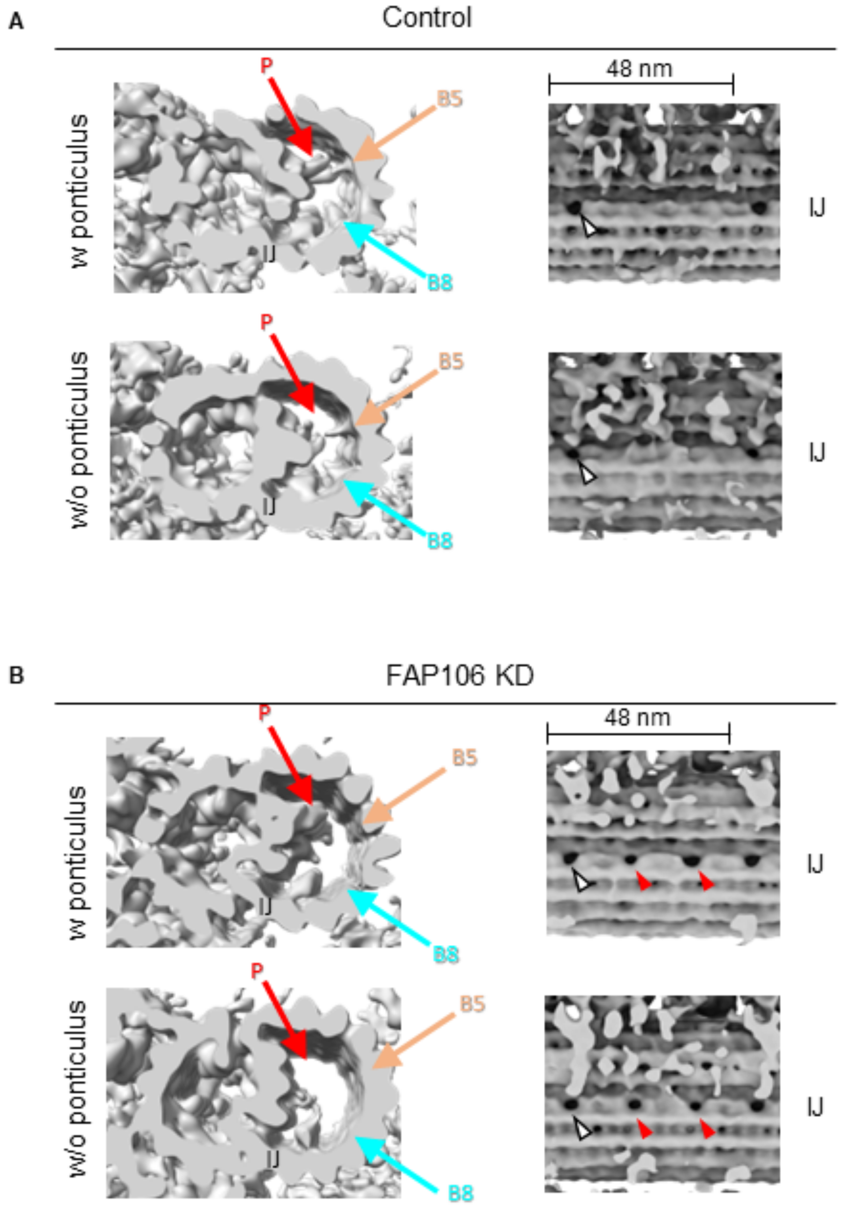


**Figure S9. Sub-tomogram averages from single flagella.** Sub-tomogram averaging was performed using particles from individual flagella in control (A) or FAP106-KD (B) samples. Each averaged tomogram shows the average obtained with particles from only a single flagellum. Individual flagella with (w) or without (w/o) clear ponticulus densities (red arrows) were identified as examples of old and new flagella, respectively, in both samples. White arrowheads show the IJ hole found in control samples. Loss of MIP B5 (peach arrow), MIP B8 (teal arrow), and appearance of extra IJ holes (red arrowheads) are dependent upon loss of FAP106, and do not show age-dependent differences. Number of particle images used for each averaged structure are listed in Supp. Table S1.**Supplemental Table S1. CryoET data collection statistics**

| | Control dataset 1 | FAP106 KD | MC8 KD | Control dataset 2 |
|---|---|---|---|---|
| **Data collection and processing** | | | | |
| Microscope | Titan Krios | Titan Krios | Titan Krios | Titan Krios |
| Magnification | x53,000 | x53,000 | x53,000 | x53,000 |
| Voltage (kV) | 300 | 300 | 300 | 300 |
| Total Electron exposure (e–/Å2) | 110 | 110 | 110 | 110 |
| Slit width (eV) | 20 | 20 | 20 | 20 |
| Detector | K3 | K3 | K3 | K3 |
| Defocus range (µm) | -3.5 to -5.5 | -3.5 to -5.5 | -3.5 to -5.5 | -3.5 to -5.5 |
| Pixel size (Å) | 1.69 | 1.69 | 1.69 | 1.69 |
| Tilt-series increment | ±2° | ±2° | ±2° | ±2° |
| Tilt-series scheme | dose-symmetric* | dose-symmetric* | dose-symmetric* | dose-symmetric* |
| Tilt-series range | ±60° | ±60° | ±60° | ±60° |
| Tilt-series collected | 52 | 51 | 69 | 17 |
| Tilt-series used | 20 | 18 | 52 | 9 |
| **Data processing** | | | | |
| Software tilt-series alignment | IMOD# | IMOD# | IMOD# | IMOD# |
| Software final reconstruction | PEET& | PEET& | PEET& | PEET& |
| Initial particle images (no.) | 1616 | 1014 | 2626 | 626 |
| Final particle images (no.) | 1014 | 728 | 1936 | 538 |
| # of particle images used for structure with ponticulus in Fig. S9 | 72 | 50 | na | na |
| # of particle images used for structure without ponticulus in Fig. S9 | 64 | 60 | na | na |
| Pixel size final reconstruction (Å) | 6.76 | 6.76 | 6.76 | 6.76 |
| Map resolution (Å) | 18.5 | 19.5 | 16.5 | 24 |
| FSC threshold | 0.143 | 0.143 | 0.143 | 0.143 |

*Dose-symmetrical tilt-scheme as described in (*7*)

#IMOD (*8*)

&PEET (*9*)

**Supplemental Table S2: Primers**

| | | Restriction Site | Primer Sequence |
|---|---|---|---|
| FAP106 (KD) | Forward | XbaI | GCTCTAGACCAGCAAAGAAACCAACGGG |
| FAP106 (KD) | Reverse | HindIII | CCCAAGCTTACTGCATTCAGCTCCTTCCC |
| FAP106 (qRT PCR) | Forward | - | ACCTGATCGAGCCAGATGTG |
| FAP106 (qRT PCR | Reverse | - | ATTCTCGCCGCCTACGTTTG |
| RPS23 (qRT PCR) (*10*) | Forward | - | AGATTGGCGTTGGAGCGAAA |
| RPS23 (qRT PCR) (*10*) | Reverse | - | GACCGAAACCAGAGACCAGCA |
| MC3 (downstream sgRNA) | Forward | - | gaaattaatacgactcactataggGTCATCTGACTTACACTGATgttttagagctagaaatagc |
| G00 (sgRNA) (*11*) | Reverse | - | aaaagcaccgactcggtgccactttttcaagttgataacggactagccttattttaacttgctatttctagctctaaaac |
| MC3 (NG repair template) | Forward | - | CACTTTTGTCATCGAATCGTATATGACTACcttGTCTCGAAAGGTGAGGA |
| MC3 (NG repair template) | Reverse | - | ATACGTCAACCATATCCTTTTGTCTCAGTGccaatttgagagacctgtgc |
| MC3 (KD) | Forward | XbaI | GCTCTAGAAAAAACAGAACGTGACGCGG |
| MC3 (KD) | Reverse | HindIII | CCCAAGCTTTATTCGTTGGGCTGTGCTGT |
| MC5 (NG) | Forward | - | ACTTACACGCAGTGGCGTCTAACGATACCTACAGGCCGGGAGTCCACAACGTTAGCAAGTTTGGGGAAACATCGAGCAAGGGTTCTGGTAGTGGTTCCGG |
| MC5 (NG) | Reverse | - | CAATCCCCTCCTCTCTACTGTTTCTGTACATTTGCTTCCTCTGGTGATTGGCGATCATCTAGTGATGTTGTTATTTGAAGCCAATTTGAGAGACCTGTGC |
| MC5 (KD) | Forward | XbaI | GCTCTAGACACAAGCCAAAGTTCCACGG |
| MC5 (KD) | Reverse | HindIII | CCCAAGCTTTGCTAACGTTGTGGACTCCC |
| MC8 (NG) | Forward | - | TATATGTGGCGGGTCGGAAAGAAAACCGACGAGCCGCAACACTATCAAAGAATAGAGTTATAGCAACTTTTTATGGTGACCTTGTCTCGAAAGGTGAGGA |

| | | | |
|---|---|---|---|
| MC8 (NG) | Reverse | - | ACTTGCTGGAGGCAGCTTTATCCACACACAACCGCACAGACTTTCCTCTTGTGTTCTCTGTCCCAAATCTCACCCACTCACCAATTTGAGAGACCTGTGC |
| MC8 (KD) | Forward | XbaI | GCTCTAGAAACTTGCAACGCAGCATCAG |
| MC8 (KD) | Reverse | HindIII | CCCAAGCTTTAACCAAGTACTGCCAGCGG |
| MC15 (NG) | Forward | - | CAAAAGGTGCGACTGGTGTCTGGAAACCCAATACATTTGAATGCACAAGTGTAGTTCAATCATGTTTGCGCCGTTTTTATCTTGTCTCGAAAGGTGAGGA |
| MC15 (NG) | Reverse | - | AGCAAAAACTACGCACGCAGCCGCCCATTTGAAGCGCTACGCAACCTCCTCTTGCCTTATTCTAAGACATTAGGCTCTACCCAATTTGAGAGACCTGTGC |
| MC15 (KD) | Forward | XbaI | GCTCTAGAGGACACCATTCCGTCTCGTT |
| MC15 (KD) | Reverse | HindIII | CCCAAGCTTGAGGTTTAGGAGGCGCAAGT |
| FAP45 (APEX) | Forward | - | GAAGGAGTTAGAGGAACTCGGTGTGCCTGAAGAGTACTGCCAGGCACTGCAGAAGAAGATGAAGGTCAAGGTGGCCCGGCGAGGTTCTGGTAGTGGTTCC |
| FAP45 (APEX) | Reverse | - | AGGGAAAAAAGAGTGACTAGGGCAATACCAAAAATATAATGCGCGTATTAATACAAAAGTTCCTGCGGTTAAAAATGTAACCAATTTGAGAGACCTGTGC |
| FAP52 (APEX) | Forward | - | ACGGCAAGAAAATTGTGTCCGTTGGTGACGAAGGCGCTATTATGATTTGGTCTGTCTGTGACTTGGAGTTTAAGACGCTGGGTTCTGGTAGTGGTTCC |
| FAP52 (APEX) | Reverse | - | ACACACATACACACACACATACATATATTTATAATATAGAGCGTCAAAGGGGAGGAGTGCACGCACATAGATAAATAGGACCAATTTGAGAGACCTGTGC |

#### Supplemental Methods

##### APEX2-dependent proximity labeling and purification of biotinylated proteins

APEX proximity labeling identifies proteins in proximity to an APEX-tagged protein, "the bait", based on biotinylation and subsequent streptavidin purification and proteomic identification (*6, 12*). We used as bait, two MIP proteins known to be inside the B-tubule of the DMT, FAP45 and FAP52, and one axonemal protein known to be outside the DMT, DRC1 (Supp. Fig. S6). APEX2-dependent biotinylation was performed as described (*6*) with the following modifications. For fluorescence microscopy, cells were biotinylated as described, then fixed with 0.2% paraformaldehyde. Cover slips were blocked in Dulbecco's phosphate buffered saline (DPBS) + 8% normal donkey serum (NDS) + 2% BSA for 1 h at room temperature before incubation with 1:1000 anti-PFR primary antibody (*13*) overnight at 4°C. 1:200 streptavidin-Alexa 594 was added during the incubation with 1:1500 donkey anti-rabbit Alexa 488 secondary antibody. After post-fixation of coverslips with 4% PFA, coverslips were washed three times in DPBS. Images were acquired on a Zeiss Axioskop II microscope with Axiovision software and processed with Zen software. For shotgun proteomics, 2e8 cells were resuspended at 2e7 cells/mL in growth medium supplemented with 5 mM biotin-phenol for 1h. Cells were treated with 50 µM $H_2O_2$ for 1 min before quenching. After two washes with quenching solution, cells were washed with DPBS and resuspended at 2e8 cells/mL in Lysis buffer (PEME buffer + 1% NP40 + 4x Sigmafast EDTA-free protease inhibitor cocktail). 40 units of Turbo DNase (Thermo Fisher Scientific) were added and the samples were incubated at room temperature for 15 min. After centrifugation, the detergent-insoluble pellet was washed once in Lysis buffer and boiled for 5 min in Lysis buffer + 1% SDS. Purification of biotinylated proteins with streptavidin beads was performed as described (*6*). Two independent biological replicates were prepared and analyzed by shotgun proteomics for each target protein (FAP45-APEX, FAP52-APEX, DRC1-APEX).

##### Shotgun proteomics

*Sample Digestion and Desalting*: The proteins bound to streptavidin beads were resuspended in digestion buffer (8M urea, 0.1 M Tris, pH 8.5) and reduced and alkylated via sequential 20-minute incubations of 5mM TCEP 10mM iodoacetamide at room temperature in the dark while being mixed at 1200 rpm in an Eppendorf thermomixer. Proteins were then digested by the addition of 0.1µg Lys-C (FUJIFILM Wako Pure Chemical Corporation, 125-05061) and 0.8µg Trypsin (Thermo Scientific, 90057) while shaking at 37°C overnight. The digestions were quenched via addition of formic acid to a final concentration of 5% (v/v). Each sample was desalted via C18 tips (Thermo Scientific, 87784) and then resuspended in 5% formic acid before analysis by LC-MS/MS.

*LC-MS Acquisition and Analysis*: Peptide samples were separated on a 75uM ID x 25cm C18 column packed with 1.9µm C18 particles (Dr. Maisch GmbH) using a 140-minute gradient of increasing acetonitrile and eluted directly into a Thermo Orbitrap Fusion Lumos mass spectrometer where MS/MS spectra were acquired by Data Dependent Acquisition (DDA). Database search was performed by using ProLuCID (*14*) and DTASelect2 (*15, 16*) implemented in Integrated Proteomics Pipeline IP2 (Integrated Proteomics Applications) and searched against a user assembled database consisting of all protein entries from the TriTrypDB (*17*) for *T. brucei* strain 927 (version 7.0). A PSM-level false positive rate cutoff was set at 1% as estimated by a target-decoy database competition strategy, protein and peptide identifications were filtered by DTASelect2 and a minimum of two unique peptides per protein were required for the confident protein identification. Raw proteomic data have been deposited to the Mass Spectrometry Interactive Virtual Environment (MassIVE) under accession ID MSV000090660.

**Filtering criteria to identify MIP candidates (MCs)**

For comparison of proteins identified in MIP (FAP45-APEX and FAP52-APEX), and non-MIP control (DRC1-APEX) samples, proteomics data were parsed using the Integrated Proteomics Pipeline IP2. The output from the Protein Identification STAT Compare tool (IDSTAT_COMPARE) in IP2 was filtered in Excel (Supp. Dataset S2). We first compared relative abundance for proteins in the MIP vs DRC1 bait samples. The threshold abundance ratio for inclusion as a MIP candidate is based on the average FAP45/DRC1 and FAP52/DRC1 ratios obtained for other known MIPs. Additional filters were subsequently applied to identify the most promising MIP candidates (MCs). In summary, proteins are identified as MCs based on: i) average FAP45/DRC1 and FAP52/DRC1 ratio meeting MIP threshold in APEX2 proteomics experiments, ii) similar abundance to known MIPs in APEX2 proteomics experiments, iii) localization to the flagellum from published data (*18, 19*), iv) absence of prior functional annotation, and v) lack of homologs outside kinetoplastids. The exact filtering criteria are as follows and in Supplementary Dataset S2.

i. Average FAP45/DRC1 and FAP52/DRC1 ratio meeting MIP threshold in APEX2 proteomics experiments: Average of FAP45/DRC1 and FAP52/DRC1 abundance ratios (Avg MIP/DRC1 ratio) $\geq$ 1.235, where 1.24 is the lowest average abundance ratio observed for known B-tubule MIPs, considering PACRGs as B-tubule MIPs based on their exposure to the B-tubule lumen, but excluding FAP20, which for unknown reasons did not show enrichment in MIP samples relative to the non-MIP control.
ii. Relative abundance comparable to known B-tubule MIPs in APEX2 proteomics experiments: Average abundance in FAP45-APEX samples $\geq$ 0.00126 and average abundance in FAP52-APEX samples $\geq$ 9.59e-4, where 0.0013 is the lowest average abundance observed for known B-tubule MIPs (as defined in i.) in the FAP45-APEX samples and 9.59e-4 is the lowest average abundance observed for known B-tubule MIPs (as defined in i.) in the FAP52-APEX samples.
iii. Flagellum localization according to Tryptag (*18, 19*)
iv. No homolog in *Chlamydomonas* or *Tetrahymena* based on OrthoMCL and/or reciprocal BLAST search
v. Not previously characterized (annotated as hypothetical or domain of unknown function)

Based on these criteria, we identified 15 MIP Candidates (MCs), named in no particular order (Supp. Dataset S2).

**Fluorescence microscopy**

Detergent-extracted cytoskeletons were prepared on cover slips as described for measurement of flagellum length. Cover slips containing cytoskeletons were rinsed in PEME and mounted onto slides for NeonGreen (NG) fluorescence microscopy. Immunofluorescence microscopy of biotinylation in whole cells was performed as described in (*6*). Images were acquired on a Zeiss Axioskop II fluorescence microscope with a Plan-Apochromat 100x/1.4 objective lens and Axiovision software or on a Zeiss Axio Imager Z1 fluorescence microscope with a Plan-Apochromat 63x/1.4 objective lens and Zen software.

#### Supplementary Datasets

##### Supplementary Dataset S1

TMT proteomics data comparing detergent-extracted flagellum skeletons. Each experiment is shown on a separate tab: FAP106 KD vs 29-13 (Ctrl); MC3 KD vs MC3-NG; MC5 KD vs MC5-NG; MC8 KD vs MC8-NG; MC15 KD vs MC15-NG.

##### Supplementary Dataset S2

APEX2-based proximity proteomics data and analysis. ‘Unfiltered data’ tab contains the full list of proteins identified. Columns in ‘Filtered data’ tab shows filters used for assigning MIP candidates (MC1-15). ‘MC List’ tab lists MC1-15 gene IDs and Avg MIPs/DRC1 ratios.
